# Only females show a stable association between neuroticism and microstructural asymmetry of the cingulum across childhood and adolescence: A longitudinal DTI study

**DOI:** 10.1101/2021.08.31.458188

**Authors:** Anna Plachti, William FC Baaré, Louise Baruël Johansen, Wesley K. Thompson, Hartwig R. Siebner, Kathrine Skak Madsen

## Abstract

Neuroticism is characterized by a tendency to experience negative and anxious emotions. This personality trait is linked to an increased risk of anxiety and mood disorders. In a cross-sectional 3T diffusion tensor imaging (DTI) study in children and adolescents, we found an association between neuroticism and a relative imbalance between left and right (i.e., asymmetry) fractional anisotropy (FA) in the cingulum and white matter underlying the ventromedial prefrontal cortex with opposite directions in females and males. Here we analyzed the longitudinal follow-up DTI data, which was acquired in 76 typically-developing 7- to 18-year-olds, including up to 11 scans per subject. Neuroticism was assessed up to four times. Our longitudinal DTI measurements substantiate robust associations between higher neuroticism scores and increased left relative to right cingulum FA in females and decreased left relative to right cingulum FA in males. In females, the association was already present in late childhood and with a stable expression across childhood and adolescence. In males, the association gradually emerged during adolescence. Future longitudinal studies should clarify which neurobiological factors (e.g., genetic variation, prenatal stress, sex hormones) contribute to the sex-specific associations in the relationship between neuroticism and interhemispheric microstructural asymmetry of the cingulum.

**Highlight:**

- We analyzed a unique longitudinal DTI dataset covering late childhood and adolescence.
- In the cingulum, left-right fractional anisotropy (FA) asymmetry scaled with neuroticism.
- Females displayed a stable association of neuroticism with increased cingulum asymmetry.
- Males showed an association between neuroticism and decreased cingulum FA asymmetry.
- The association in males became more accentuated during adolescence

## 1. Introduction

Adolescence is associated with an increased incidence of neuropsychiatric disorders including mood disorders such as anxiety or depression (Bradley, 2001; Paus et al., 2008). The risk of developing mood disorders is increased in individuals scoring high on the personality trait neuroticism (Hettema et al., 2006; Kendler & Myers, 2010; Tully et al., 2015). Neuroticism reflects an individual’s tendency towards experiencing negative emotions, such as anxiousness, sadness, worry, and difficulties coping with stress (Bienvenu et al., 2001). Neuroticism is moderately heritable (Sanchez-Roige et al., 2018; van den Berg et al., 2014) and the association between neuroticism and anxiety/depressive symptoms and major depression seems largely to be due to common genetic risk factors (Kendler & Myers, 2010; Luciano et al., 2018). Moreover, females generally score higher on neuroticism than males throughout most of the lifespan (Chapman et al., 2007; Schmitt et al., 2017). This difference may already become apparent in adolescence around the age of 14 years (De Bolle et al., 2015), consonant with evidence that an equal female-male prevalence of anxiety and mood disorders before puberty changes to a 2:1 female-male prevalence after puberty (Goodwin & Gotlib, 2004; Paus et al., 2008).

Neuroimaging studies investigating the neural correlates of neuroticism and other negative emotionality traits such as harm avoidance have mostly been conducted in adult cohorts (Mincic, 2015; Opel et al., 2020; Privado et al., 2017; Servaas et al., 2013). Structural magnetic resonance imaging (MRI) studies have shown associations between higher neuroticism scores and a wide range of brain regions, including smaller total brain volume and smaller fronto-temporal surface area (Bjørnebekk et al., 2013), thicker left inferior parietal cortex (Privado et al., 2017), larger left amygdala and anterior parahippocampal gyrus volumes and smaller grey matter volumes in anterior brain regions such as orbitofrontal and cingulate cortex with a predominance in the left hemisphere (Mincic, 2015). Diffusion-weighted imaging (DWI) studies have revealed associations of neuroticism and other negative emotionality-related personality traits with global white matter tissue measures of fractional anisotropy (FA) and mean- and radial diffusivities (MD, RD) (Bjørnebekk et al., 2013), as well as with reduced FA in major white matter fiber tracts such as uncinate fasciculus and cingulum bundle (Mincic, 2015; Privado et al., 2017). Despite the distributed observed associations, many studies have implicated a fronto-limbic network in negative emotionality traits, including the cingulum, uncinate fasciculus and the white matter underlying the ventromedial prefrontal cortex (vmPFCWM) (Cremers et al., 2010; Forbes et al., 2014; Gardiner et al., 2015; Madsen et al., 2012; Madsen et al., 2018; Mincic, 2015; Moshirian Farahi et al., 2019).

Recently, in a cross-sectional study of the same sample of children and adolescents aged 10-15 years included in the present study, we found that neuroticism was associated with the relative balance between left and right hemispheric white matter tracts, rather than the absolute FA values of these white matter tracts. Higher neuroticism scores were associated with decreased left relative to right cingulum FA in males, while in females, higher neuroticism scores were related to increased left relative to right cingulum and ventromedial prefrontal white matter FA. Our finding in adolescent males aligned well with our similar observation in a cohort of healthy adults aged 19-86 years that contained approximately twice as many males as females (Madsen et al., 2012). Furthermore, the observed associations in children and adolescents became stronger with increasing age in males, but not in females (Madsen et al., 2018). Although intriguing, the latter needs to be scrutinized using a longitudinal design. Notably, the cingulum and uncinate fasciculus undergo continuous maturation, i.e., age-related increases in FA and decreases in MD, throughout childhood and adolescence and into early adulthood (Lebel & Beaulieu, 2011). Finally, whereas sex differences in associations between neuroticism and brain structure may already be present in adolescence (Blankstein et al., 2009; Madsen et al., 2018), the possible effect of sex on negative emotionality-related traits and brain structure has not been systematically investigated yet and findings are generally inconclusive (Avinun et al., 2020; Mincic, 2015; Nostro et al., 2017).

In the present longitudinal study, we aimed to elucidate sex differences in fronto-limbic white matter correlates of neuroticism and to characterize how the relationship between neuroticism and fronto-limbic white matter might change with age across childhood and adolescence. To this end, we investigated typically-developing children and adolescents aged 7 and 18 years, who underwent DWI up to 11 times and were assessed on neuroticism up to four times. Based on our cross-sectional findings in the same sample (Madsen et al., 2018), we expected that our longitudinal data would affirm the previously observed cross-sectional associations between neuroticism and cingulum and vmPFCWM FA asymmetries. Specifically, we expected that higher neuroticism scores would be associated with increased left relative to right cingulum and vmPFCWM FA asymmetries in females, and to decreased left relative to right cingulum FA asymmetry. Furthermore, we expected that, the neuroticism associations with cingulum and vmPFCWM FA asymmetry would differ as a function of age within the investigated age range.

## 2. Material and methods

### 2.1 Study design and participants

The present longitudinal study included data from 76 typically-developing children and adolescents (47 females, 29 males) aged 7.5-18.9 years (mean = 12.4, standard deviation (SD) = 2.4), who were enrolled in the longitudinal HUBU (“*Hjernens Udvikling hos Børn og Unge*”) study. The HUBU study was initiated in 2007, where 95 participants (55 females, 40 males) aged 7-13 years and their families were recruited from three elementary schools in the Copenhagen suburban area. All children whose families volunteered were included, except for children who according to parent-reported screening questionnaires had any known history of neurological or psychiatric disorders or significant brain injury. Prior to participation, children and their parents were informed about the aims and procedures of the study in oral and written language and in accordance with children’s assent, all parents gave their informed written consent. Further, informed written consent was obtained from the participants themselves when they turned 18 years. The study was approved by the Ethical Committees of the Capital Region of Denmark (H-KF-01-131/03 and H-3-2013-037) and performed in accordance with the Declaration of Helsinki. Participants were assessed up to 13 times with six months intervals between the first 10 sessions, one-year interval for the 11^th^ assessment, and three-year intervals for assessments 12 and 13.

The present study included data from the first 11 assessments of the HUBU study. A total of 19 participants (8 females, 11 males) were excluded entirely from the present longitudinal study due to incidental findings, psychiatric diagnoses or lack of neuroticism data (see Supplement methods). A total of 675 valid observations (range = 3-11, mean = 8.8 observations per subject) were included in the statistical analyses. A summary of the longitudinal assessments and participants’ age at each assessment is presented in Figure 1. According to the Edinburg Handedness Inventory (EHI), 68 participants were right-handed (EHI score ≥40) and 8 were left-handed (EHI score ≤ -40). Highest level of maternal and paternal education was transformed into years of education using national norms and used to estimate average years of parental education for all participants (mean = 13.8, SD = 2.0). Data from the HUBU cohort has previously been used in cross-sectional (Angstmann et al., 2016; Gonzalez et al., 2021; Klarborg et al., 2013; Madsen et al., 2011; Madsen et al., 2010; Madsen et al., 2018; Vestergaard et al., 2011) and longitudinal (Madsen et al., 2020) studies examining brain-behavioral relationships.

**Figure 1.**
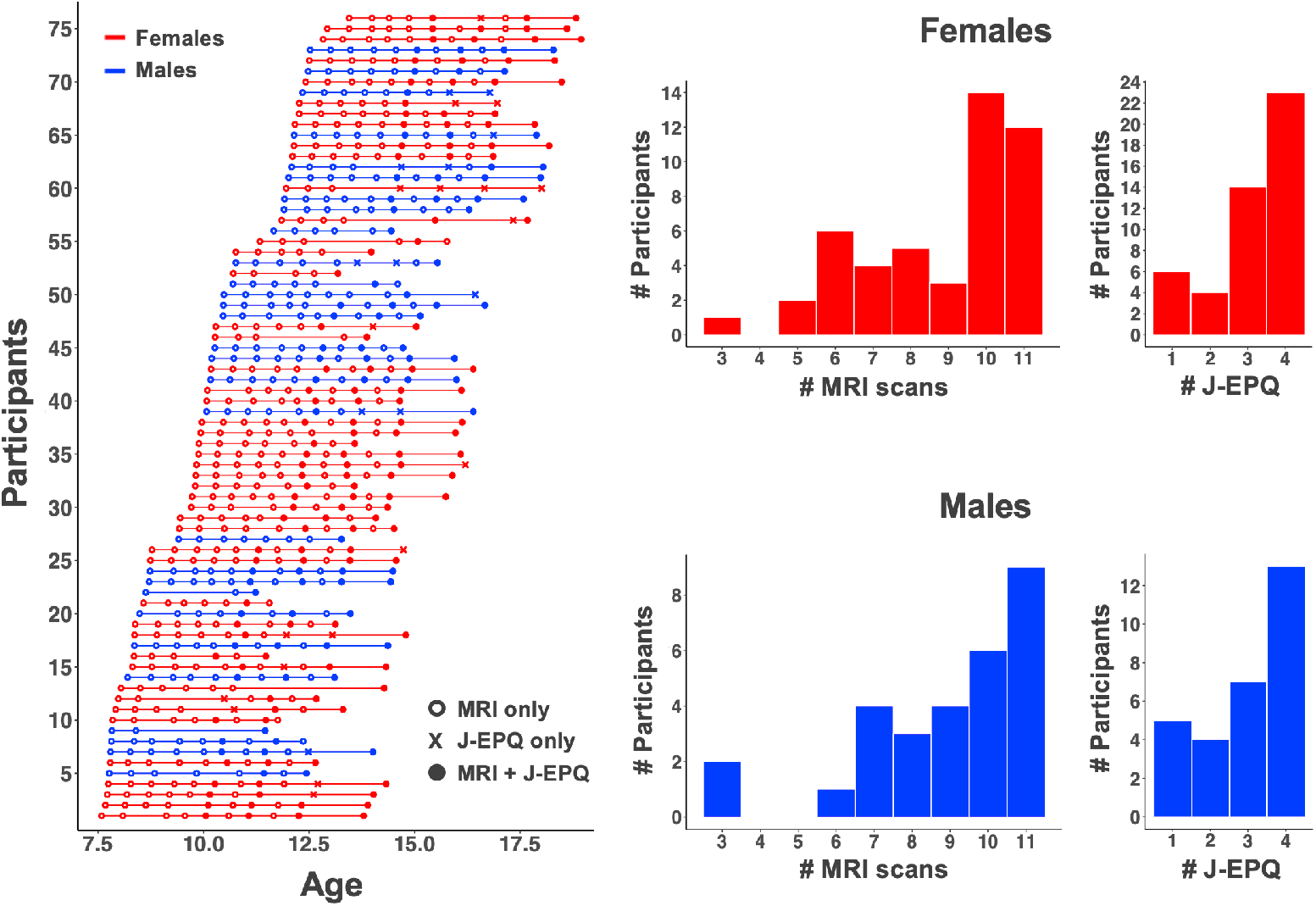
Left: Number of longitudinal observations and age at each assessment for each of the 76 participants. Assessments including MRI and the Junior Eysenck Personality Questionnaire (J-EPQ, in the present study only neuroticism scores were used - see text) are shown as closed circles, MRI only as open circles, and J-EPQ only as “x”. Right: Histograms depicting the number of participants that have a certain number of MRI scans or J-EPQ assessments, for males and females separately. Females (n=47) are shown in red, and males (n=29) in blue. # = “number of”

### 2.2 Personality assessment

Participants completed an adapted Danish version of the 81-item Junior Eysenck Personality Questionnaire (J-EPQ) (Eysenck & Eysenck, 1975; Nyborg et al., 1982) on the same day as the MR-scanning in rounds 6, 8, 10 and 11. The J-EPQ measures three major dimensions of personality, i.e., neuroticism, extraversion and psychoticism. Individual J-EPQ statements were read out loud by the test administrator and participants subsequently rated how well each statement fitted them. In HUBU, we extended the original “Yes” or “No” rating scale to: Strongly agree / Agree / Disagree / Strongly disagree, to allow for a more fine-grained evaluation of the personality traits. The answers were scored on a 0-3-point scale, with 0 reflecting strongly disagree. In the present study, we focused on the neuroticism scale that consists of 20 items. The neuroticism scale showed good to excellent internal consistency using either the 2-point or 4-point rating scales (Madsen et al., 2018).

### 2.3 Image acquisition

All participants were scanned using a 3T Siemens Magnetom Trio MR scanner (Siemens, Erlangen, Germany) with an eight-channel head coil (Invivo, FL, USA). T1-weighted and T2-weighted images of the whole head were acquired using 3D MPRAGE and 3D turbo spin echo sequences. Whole brain diffusion-weighted (DW) images were acquired using a twice-refocused balanced spin echo sequence that minimized eddy current distortion (Reese et al., 2003) with 10 non-DW and 61 DW images (b = 1200 s/mm^2^). A gradient echo field map was acquired to correct B0 field distortions. Sequence parameters are given in the Supplementary methods.

### 2.4 Image evaluation

An experienced neuroradiologist evaluated all baseline MRI scans, and all, but one, were deemed without significant clinical pathology. Prior to analysis and blind to behavioral data, all raw MR-images were visually inspected to ascertain image quality and excluded if not of sufficient quality (see Supplementary methods).

### 2.5 Construction of the diffusion tensor images

“Image preprocessing was done using MATLAB scripts that were mainly based on SPM 8 routines (Wellcome Department of Cognitive Neurology, University College London, UK). The T1- and T2-weighted images were rigidly oriented to MNI space (six-parameter mutual information) and corrected for spatial distortions due to nonlinearity in the gradient system of the scanner (Jovicich et al., 2006) (note that at this point no reslicing was performed). T2-weighted images were then rigidly co-registered to the T1-weighted image. To align the DWI images to the T1-weighted image, the mean b0 image was first rigidly registered to the T2-weighted image, after which all DW images were co-registered to the mean b0 image (no reslicing). Next, all co-registered DWI images were corrected for spatial distortions using a voxel displacement map based on the acquired b0 field map (Andersson et al., 2001) and the scanner-specific gradient non-linearity profile (Jovicich et al., 2006). Subsequently, all images were resliced using tri-linear interpolation. Importantly, the above procedure ensures that only one re-slicing step was employed. Diffusion gradient orientations were adjusted to account for any applied rotations. The least-squares-fit by non-linear optimization, employing a Levenburg-Marquardt algorithm and constrained to be positive definite by fitting its Cholesky decomposition, implemented in Camino was used to fit the diffusion tensor (DT) (Jones & Basser, 2004). Finally, for each subject we estimated movement during DWI scanning by calculating the root mean square deviation (RMS) of the six rigid body transformation parameters resulting from co-registering DWI images to the mean b0 image (Jenkinson & Smith, 2001; Taylor et al., 2016), using a spherical volume radius of 60 mm to approximate the brain.” (Madsen et al., 2020).

### 2.6 Spatial normalization of the longitudinal diffusion tensor images

“The diffusion tensor images were spatially normalized using DTI-TK, which uses the high dimensional information of the diffusion tensor to achieve highly accurate normalizations (Zhang et al., 2007). We employed an unbiased longitudinal two-step approach (Keihaninejad et al., 2013). First, within each subject all DT images over all time points were registered together to create within-subject DT image templates. Secondly, within-subject template DT images were registered together to create a between-subject DT template image. Concatenation of the within- and between subject registration deformation fields was used to warp individual DWI volumes into a common study specific space. Fractional anisotropy (FA), axial diffusivity (AD = λ1) and radial diffusivity (RD = (λ2 + λ3) / 2) images were created. Finally, non-brain voxels in FA and diffusivity images were removed by employing a brain mask based on warped b0 images” (Madsen et al., 2020).

### 2.7 Tract-based spatial statistics

Tract-Based Spatial Statistics (TBSS) (Smith et al., 2006) was used to create a mean FA skeleton, representing the centers of all tracts common to the group, by first aligning the between-subject FA template image (from DTI-TK) to MNI space using affine registration (flirt, FSL), and then aligning the individual FA images to the template image in MNI space. A detailed description on the TBSS procedures can be found in the Supplementary methods.

### 2.8 Regions-of-interest

We extracted FA, AD and RD values from left- and right-sided regions-of-interest (ROIs) to test specific hypotheses and to determine the anatomical specificity of observed associations. White matter ROIs included the cingulum, the uncinate fasciculus (UF) and the white matter underlying vmPFC and were drawn manually onto the mean skeleton overlaid on the mean FA image using FSLview. The ROIs are shown in Figure 2. Descriptions of the ROI delineations can be found in the Supplementary methods.

**Figure 2.**
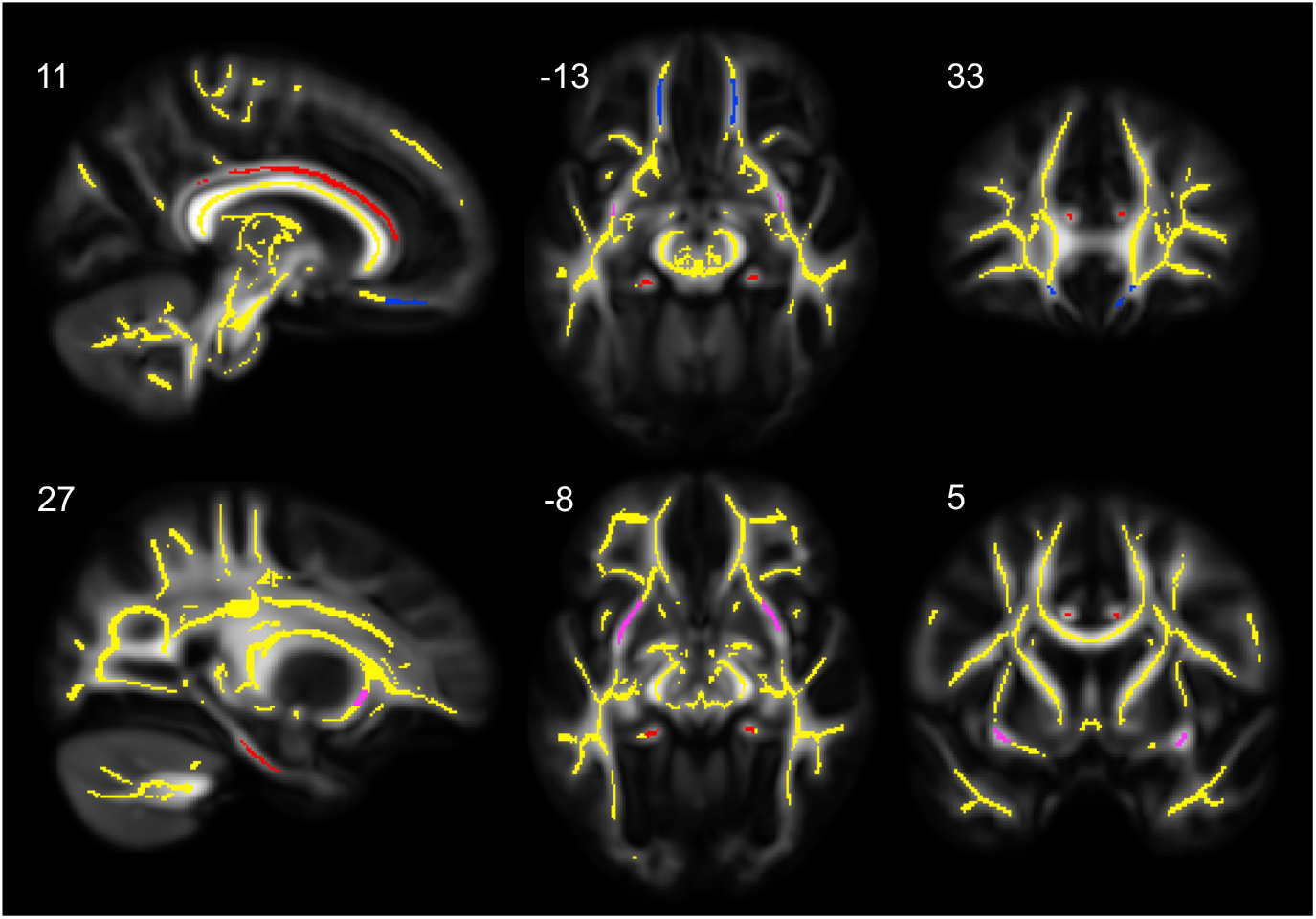
Regions-of-interest (ROIs) used in the cingulum bundle (red), uncinate fasciculus (UF, magenta) and the white matter underlying the ventromedial prefrontal cortex (vmPFC_WM_, blue) overlaid on the TBSS skeleton (yellow) and the mean fractional anisotropy (FA) map. ROIs are shown in sagittal (x), coronal (y) and axial (z) views with the corresponding MNI coordinates given above each slice.

### 2.9 Left-right asymmetry measures

Left-right ROI asymmetries were calculated for the different DTI parameters as the difference between left and right ROI values expressed as a percentage of the bilateral mean, with positive values indicating larger left relative to right measures:
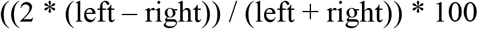

### 2.10 Statistical analyses

Statistical analyses were performed in RStudio 3.6.2 (R Core Team, 2019). The longitudinal analyses were conducted with generalized additive mixed models (GAMM) using the mgcv package, version 1.8-31 and the nlme package, version 3.1-142 (Pinheiro et al., 2013). GAMM easily handles unbalanced longitudinal data, such as unequal number of timepoints and/or unequal intervals between timepoints and uses non-linear smooth functions to fit complex longitudinal data without *a priori* assumptions of the shape of the trajectories. The *estimated degrees of freedom* (edf) denote the degrees of freedom used by the smooth function to fit the data. An edf value of one represents a linear relationship with a straight regression line, whereas increasingly larger edf values reflect increasingly more non-linear relationships.

Dependent variables were the ROI DTI or ROI DTI asymmetry measures e.g., ROI FA or ROI FA asymmetry. Sex was input as a covariate, coded with females = 1 and males = 0. The smooth terms were used for, respectively, age and age-by-sex interaction effects. Participant and within-subject age were used as random effects to fit individual intercepts and slopes, respectively. We used the default thin plate regression splines smoothing function to fit the data with default penalizing parameters. Further, we used five knots (k), representing the k-1 number of basis functions used to generate the smooth function, as we previously have shown that this reduces the risk of overfitting (Madsen et al., 2020). The smoothing parameters were estimated with the restricted maximum likelihood method (REML). All our continuous variables, i.e., age, neuroticism scores, parental education, RMS and ROI DTI measures, were z-transformed.

#### 2.10.1 Age, sex and age-by-sex effects on longitudinal neuroticism scores

Prior to testing our hypothesis, we tested for age, sex, and age-by-sex interaction effects on the longitudinal neuroticism data (neuroticism_long_) (Model 1) and possible additional handedness and parental education effects (Model 2), to assess whether neuroticism changed across age and was associated with the additional covariates.

Model 1: Neuroticism_Long_ = age + sex + age-by-sex

Model 2: Neuroticism_Long_ = age + sex + age-by-sex + parental education + handedness

#### 2.10.2 Age, sex and age-by-sex and subject movement effects on ROI FA measures

Next, we tested for age, sex, age-by-sex and RMS effects on left or right ROI FA, or ROI FA asymmetry (Model 3) and possible additional handedness and parental education effects (Model 4).

Model 3: ROI FA or FA asymmetry = age + sex + age-by-sex + RMS

Model 4: ROI FA or FA asymmetry = age + sex + age-by-sex + RMS + parental education + handedness

#### 2.10.3 Testing of a priori hypotheses: Associations between ROI FA asymmetry and neuroticism

Since neuroticism_long_ scores did not significantly change over time (see Table 1 and Figure 3), we used intra-individual mean neuroticism scores (neuroticism_mean_) when testing our hypotheses. This allowed us to include MRI scans from all time points. To test our hypotheses, neuroticism_mean_ and neuroticism_mean_-by-sex were included as independent variables-of-interest, while controlling for age, sex, age-by-sex and RMS. Dependent variables were cingulum or vmPFCWM FA asymmetry (Model 5). UF FA asymmetry was tested exploratively.

Model 5: ROI FA asymmetry = age + sex + age-by-sex + RMS + neuroticism_mean_ + neuroticism_mean_-by-sex

**Table 1.**
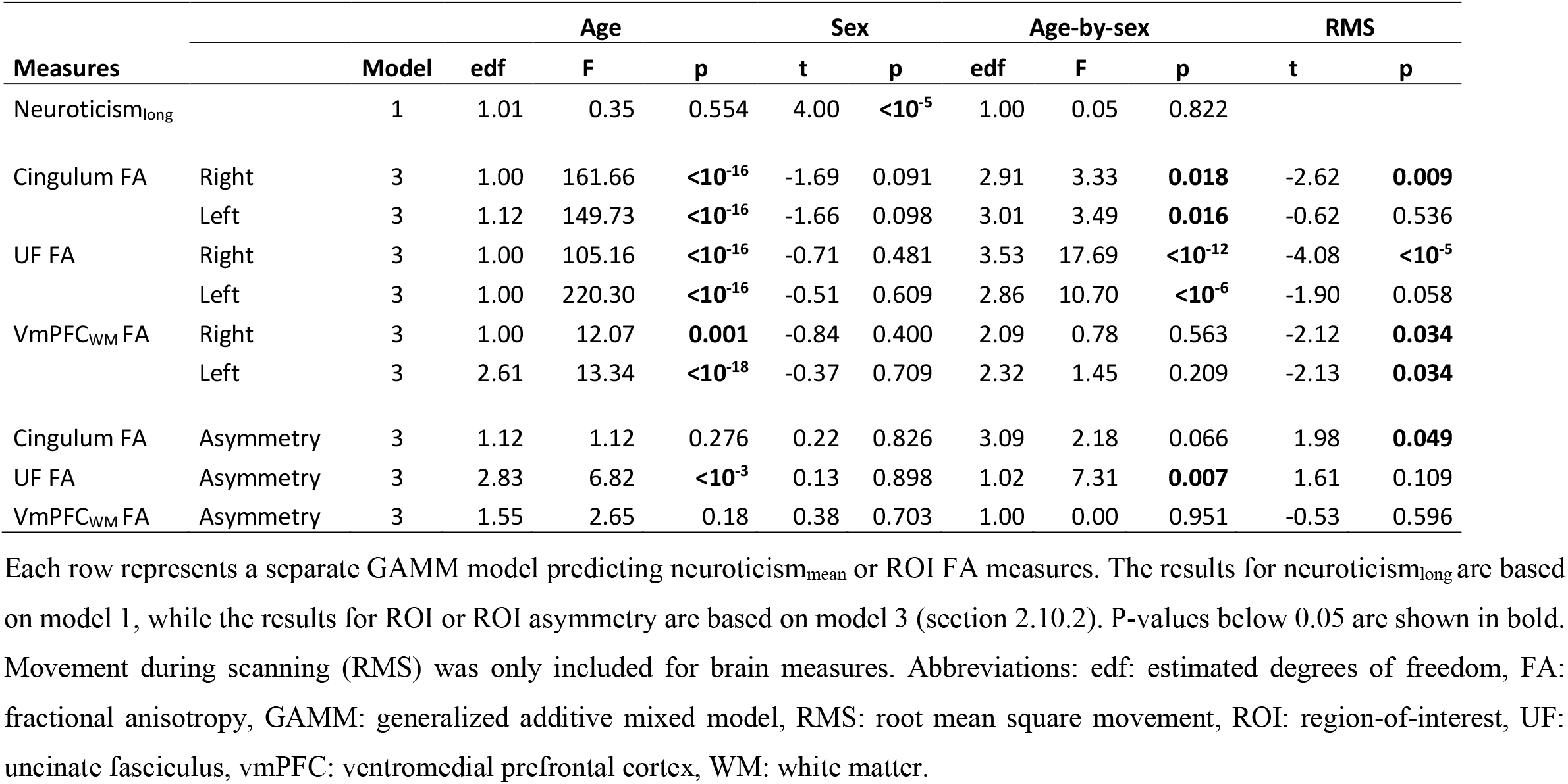
Age, sex, age-by-sex, and RMS on ROI FA, ROI FA asymmetry and neuroticism_long_.

**Figure 3.**
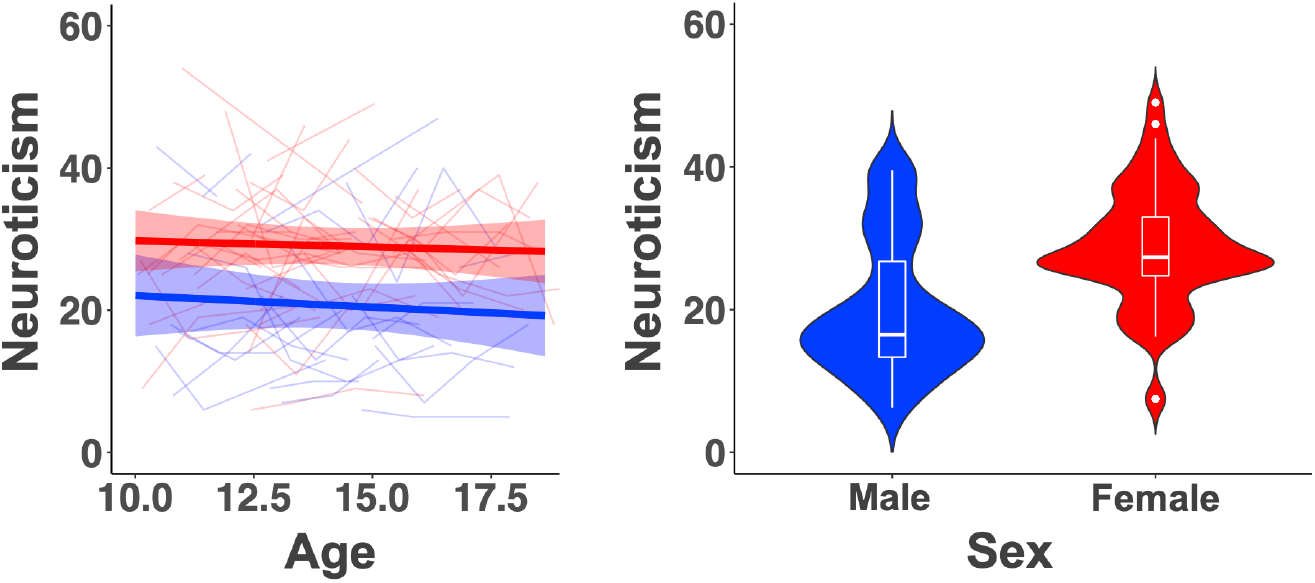
Left: Spaghetti plot of neuroticism_long_ overlaid with GAMM estimated age trajectories with shaded 95% confidence intervals for males (blue, n = 29) and females (red, n = 47). Right: Violin and box plots showing neuroticism_mean_ scores for females and males. Females had significant higher neuroticism_mean_ scores than males.

Since we were *a priori* interested in any association including neuroticism_mean_ (main and interaction effects), we applied a Bonferroni correction for four associations of interest (two for cingulum and two for vmPFC FA asymmetry). Thus, a p-value below 0.0125 was considered significant. In our exploratory analyses of UF FA asymmetry a Bonferroni corrected p-value below 0.0083 (corrected for six associations of interest, i.e., four *a priori* hypothesized, and two exploratory of UF FA asymmetry) was considered significant.

#### 2.10.4 Sex specific analyses

Contingent on observing significant neuroticism_mean_-by-sex effects on ROI FA asymmetry in Model 5, follow-up analyses were conducted in females and males separately to further investigate the neuroticism_mean_-by-sex effects (Model 6) as well as to investigate if any relationship between neuroticism_mean_ and ROI FA asymmetry in females or males changed with age (Model 7).

Model 6: ROI FA asymmetry = age + RMS + neuroticism_mean_

Model 7: ROI FA asymmetry = age + RMS + neuroticism_mean_ + neuroticism_mean_-by-age

#### 2.10.5 Follow-up analyses

Contingent on observing significant neuroticism_mean_ or neuroticism_mean_-by-age effects on ROI FA asymmetry for males and females separately, we performed several planned follow-up analyses. As no significant neuroticism_mean_-by-age effects on ROI FA asymmetry were observed (see section 3.1.1), the follow-up models were all extensions of Model 6. First, we assessed the anatomical specificity of the observed neuroticism_mean_ and ROI FA asymmetry association by including hemispheric FA asymmetry as an additional covariate. Secondly, we tested whether the observed association remained when including parental education and handedness as additional covariates. Thirdly, we assessed the contribution of absolute right and left ROI FA values to neuroticism_mean_ by analyzing them separately. Finally, to further explore the nature of our observed FA findings, we performed similar analyses as for FA on ROI AD and ROI RD asymmetries.

#### 2.10.6 Effect size maps

We generated effect size maps to provide further information about possible associations between neuroticism_mean_ and FA values across the white matter skeleton. The effect size maps display the distribution of uncorrected t-values of the associations between neuroticism_mean_ and FA in males and females separately, controlling for age and RMS (Model 6). The unthresholded t-maps have been uploaded to NeuroVault.org (Gorgolewski et al., 2015) and are available at https://neurovault.org/collections/LRMGYBHV/.

## 3. Results

### 3.1 Descriptive statistics

#### 3.1.1 Age, sex and age-by-sex effects on longitudinal neuroticism scores

There were no significant age or age-by-sex effects on the neuroticism_long_ scores (Table 1 and Figure 3), suggesting that neuroticism_long_ scores did not change significantly across childhood and adolescence. However, as expected we observed a significant effect of sex with females scoring higher than males on neuroticism_long_ (t = 4.677, p < 10^−5^) and neuroticism_mean_ (Kruskal Wallis t-test: df = 1, X^2^ = 127.47, p < 10^−16^, Figure 3). Furthermore, higher neuroticism_long_ scores were significantly associated with lower parental education (t = -2.76, p = 0.006). No significant association was observed between neuroticism_long_ and handedness (t = -1.583, p = 0.115). Since there were no significant age or age-by-sex effects on neuroticism_long_, we used the neuroticism_mean_ score of each individual for further analysis. In case a participant had only one neuroticism score (5 males, 6 females), this score was used as the participant’s ‘mean neuroticism score’.

#### 3.1.2 Age, sex, age-by-sex, and RMS effects on ROI FA and FA asymmetry

Effects of age, sex, age-by-sex and RMS on the ROI FA measures are reported in Table 1. Spaghetti plots overlaid with GAMM estimated age trajectories for males and females are shown in Figure 4. FA significantly increased with age in all right and left ROIs. Moreover, significant age-by-sex effects were found for right and left cingulum and UF FA, with males displaying more linearly increasing maturational trajectories, while females displayed more curvilinear trajectories that started to level off within the second half of the investigated age range. For ROI FA asymmetry, only the UF exhibited significant age and age-by-sex effects, with the UF changing from rightward FA asymmetry to less asymmetric FA with age in males, and slightly increased rightward FA asymmetry with age in females. No significant main effects of sex were observed for any of the ROI FA measures. Significant effects of movement during DWI (RMS) were found for several ROIs FA measures (Table 1).

**Figure 4.**
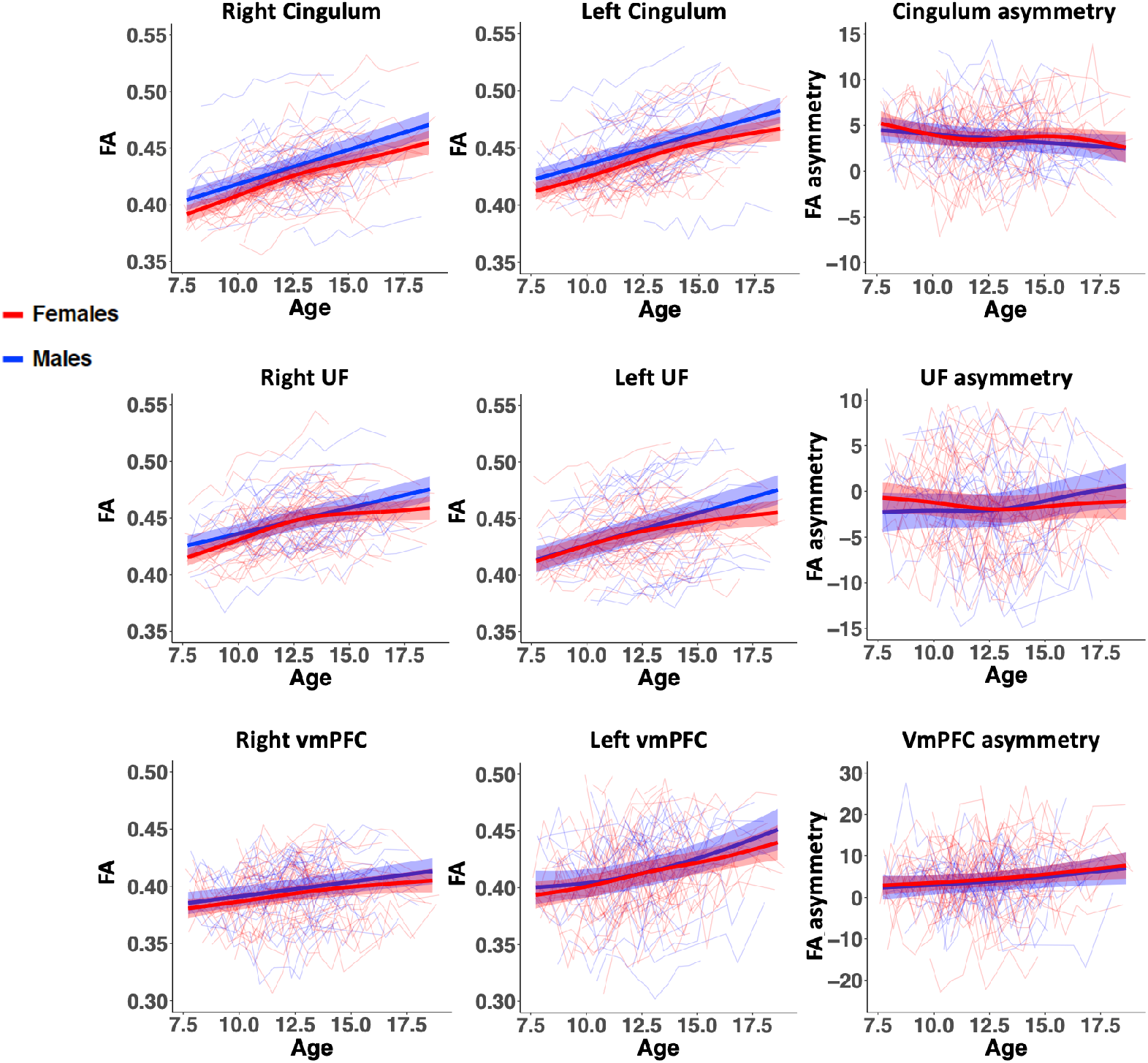
Spaghetti plots of cingulum, uncinate fasciculus (UF) and white matter underlying the ventromedial prefrontal cortex (vmPFC_WM_) fractional anisotropy (FA) overlaid with GAMM estimated age trajectories with shaded 95% confidence intervals for males (blue) and females (red).

When additionally controlling for parental education and handedness, observed age, age-by-sex and RMS effects remained, and a modest sex effect appeared, with males having higher left cingulum FA than females (t = -2.025, p = 0.043). Parental education was significantly associated with right and left vmPFCWM FA (right: t = 3.493, p = 0.0005; left: t = 2.442, p = 0.015), but not with FA in other ROIs (p’s > 0.08). Handedness was significantly associated with right UF FA (t = 2.552, p = 0.011) and right vmPFCWM FA (t = 3.399, p = 0.0007), but not with other ROI measures (p’s > 0.06).

### 3.2 Hypotheses testing

#### 3.2.1 Associations between ROI FA asymmetry and mean neuroticism

Results concerning our main hypotheses are presented in Table 2. As hypothesized, we found a significant neuroticism_mean_-by-sex interaction effect on cingulum FA asymmetry, with higher neuroticism_mean_ scores being associated with increased left relative to right cingulum FA in females and decreased left relative to right FA in males (Figure 5, left sided plot). No significant neuroticism_mean_-by-sex interaction effects were observed for vmPFCWM FA asymmetry, nor exploratory for UF FA asymmetry. Furthermore, we did not observe any significant main effects of neuroticism_mean_ on any of the ROI FA asymmetries.

**Table 2.** Associations between neuroticism_mean_ and ROI FA asymmetry

| Measures | Model | Age |  |  | Sex |  | Age-by-sex |  |  | RMS |  | Neuroticism |  | Neuroticism-by-sex |  |  |
| --- | --- | --- | --- | --- | --- | --- | --- | --- | --- | --- | --- | --- | --- | --- | --- | --- |
|  |  | edf | F | p | t | p | edf | F | p | t | p | t | p | edf | F | p |
| Cingulum FA asymmetry | 5 | 1.00 | 1.70 | 0.193 | 0.18 | 0.860 | 3.10 | 2.44 | **0.046** | 1.93 | 0.055 | -1.70 | 0.090 | 1.01 | 8.84 | **0.003** |
| VmPFC<sub>WM</sub> FA asymmetry | 5 | 1.61 | 2.56 | 0.190 | 0.06 | 0.953 | 1.00 | 0.01 | 0.920 | -0.54 | 0.587 | -0.44 | 0.662 | 1.00 | 2.28 | 0.132 |
| UF FA asymmetry | 5 | 2.80 | 6.64 | **<10<sup>-3</sup>** | 0.14 | 0.888 | 1.00 | 7.04 | **0.008** | 1.61 | 0.109 | -0.55 | 0.583 | 1.00 | 0.75 | 0.388 |

Each row represents a separate GAMM model predicting ROI FA asymmetry with neuroticism<sub>mean</sub> and neuroticism<sub>mean</sub>-by-sex, controlling for age, sex, age-by-sex and RMS. Results are based on model 5 (section 2.10.3). P-values below 0.05 are shown in bold. Abbreviations: edf: estimated degrees of freedom, FA: fractional anisotropy, GAMM: generalized additive mixed model, RMS: root mean square movement, ROI: region-of-interest, UF: uncinate fasciculus, vmPFC: ventromedial prefrontal cortex.

**Figure 5.**
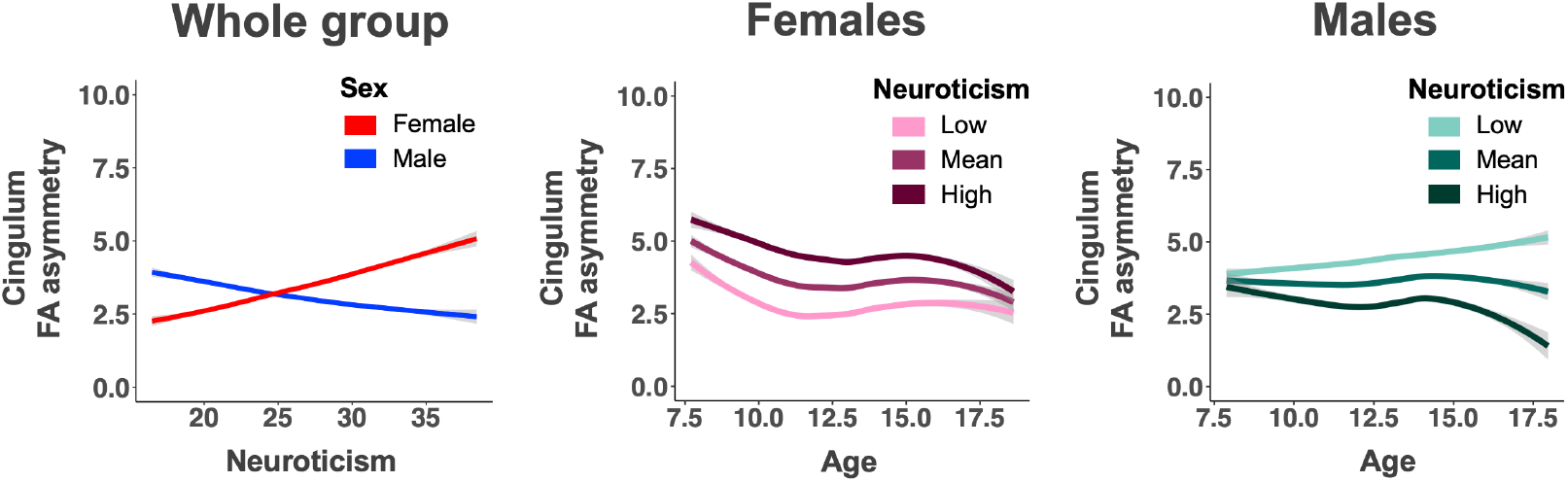
Left) Depiction of the observed significant interaction effect between sex and neuroticism_mean_ on cingulum FA asymmetry represented for age_mean_ for females (red) and males (blue), with shaded +/-95% confidence intervals. Higher neuroticism_mean_ scores were associated with increased left relative to right cingulum FA in females and decreased left relative to right cingulum FA in males. Middle and right) Depiction of the associations between neuroticism_mean_, age and cingulum FA asymmetry for respectively females (pink colors) and males (teal colors). Lines represent the predicted age trajectories for cingulum FA asymmetry, with shaded +/-95% confidence interval bands, for different levels of neuroticism: mean neuroticism is shown in medium dark shades, while +/-1 standard deviation from the mean are shown in, respectively, dark and light shades.

#### 3.2.2 Sex specific analyses

Results from the follow-up analyses of cingulum FA asymmetry in males and females separately are represented in Table 3. For females, we observed a significant effect of neuroticism_mean_, with higher neuroticism_mean_ scores being associated with higher left relative to right cingulum FA (Figure 5). In males, the association was opposite, i.e., higher neuroticism_mean_ scores were associated with lower left relative to right cingulum FA, but this association did not reach statistical significance. We did not observe significant neuroticism_mean_-by-age interaction effects. Visual inspection of the plots in Figure 5 displaying the association between neuroticism_mean_ and cingulum FA asymmetry across age, however, suggested that in females the cingulum-neuroticism_mean_ relationship might already be present in the youngest part of the included age range, while in males, an association between neuroticism_mean_ and cingulum FA asymmetry appears to become apparent in the older part of the included age range.

**Table 3.**
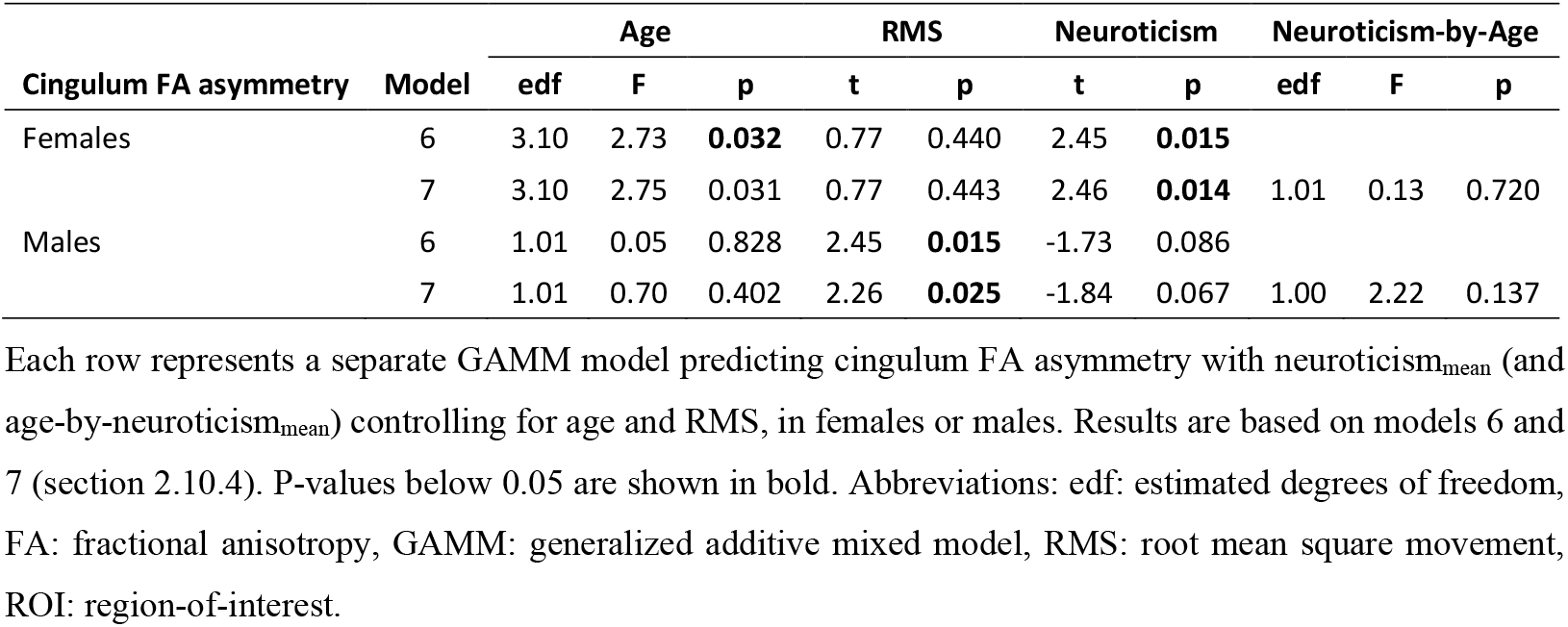
Associations of cingulum FA asymmetry and neuroticism_mean_ in females and males separately.

### 3.3 Follow-up analyses in females and males

#### 3.3.1 Follow-up analyses in females

Follow-up analyses in females showed that the association between neuroticism_mean_ and cingulum FA asymmetry remained significant (t = 2.535, p = 0.012), when hemispheric white matter FA asymmetry was included as an additional covariate in Model 6, suggesting that the relationship was not driven by individual differences in overall hemispheric asymmetry in FA. Moreover, the association between neuroticism_mean_ and cingulum FA asymmetry remained significant (t = 2.828, p = 0.005) after controlling for parental education and handedness.

Next, we assessed the contribution of left and right cingulum FA to neuroticism_mean_. Neither left nor right cingulum FA was significantly associated with neuroticism_mean_ (Left: t = - 0.248, p = 0.804; Right: t = -1.583, p = 0.114), suggesting that the association between neuroticism_mean_ and cingulum FA asymmetry was driven by the relationship between left and right cingulum FA, and not by the absolute FA values of the left or right cingulum.

Finally, to further explore the nature of our observed FA findings, we performed similar analyses as for FA (Model 6) on the cingulum AD and RD asymmetries, but no significant effects of neuroticism_mean_ on cingulum AD (t = 1.397, p = 0.163) or RD (t = -1.896, p = 0.059) asymmetry were observed.

#### 3.3.2 Post hoc analyses in males

Because the visual inspection of the plots in Figure 5 suggested that an association between neuroticism_mean_ and cingulum FA asymmetry became apparent with age in males, we performed post hoc analyses, similar to model 6, in respectively the oldest and youngest part of the age range, split by the median (12.5 years). These analyses revealed that the relationship between neuroticism_mean_ and cingulum FA asymmetry were indeed stronger in the older (t = -2.533, p = 0.013) than in the younger (t = -1.316, p = 0.191) part of the included age range.

### 3.4 Effect size maps

Effect size maps were created for males and females separately to provide further information about the distribution of the association between neuroticism_mean_ and FA across the white matter skeleton (see Supplementary results). The unthresholded t-maps can be downloaded from https://neurovault.org/collections/LRMGYBHV/.

## 4. Discussion

Our unique longitudinal DTI dataset enabled us to address two questions: Is neuroticism associated with interhemispheric microstructural asymmetry of fronto-limbic white matter tracts? If so, does this association change with age in typically-developing children and adolescents scanned up to 11 times? We found that higher neuroticism_mean_ scores were associated with increased FA in left cingulum relative to right cingulum in females, while higher neuroticism was associated with decreased FA in left relative to right cingulum in males. The association was most prominent in females, who also showed higher neuroticism scores than males. While neuroticism scores did not significantly change with age in males and females, there were apparent differences in the temporal trajectory of the association between neuroticism and FA asymmetry of the cingulum between males and females. In females the association seemed already to be present in late childhood and remain stable during adolescence, whereas the association emerged more gradually during adolescence in males. Follow-up analyses in females showed that the association was not driven by global hemispheric white matter FA asymmetry, handedness, or parental education. Moreover, the association appeared to be specific to the left-right asymmetry of FA, as it was not driven by absolute left or right cingulum FA values. Finally, we did not replicate our baseline observation of a significant link between neuroticism and FA asymmetry of vmPFCWM.

Our finding that neuroticism is associated with cingulum FA asymmetry, revealed by high temporal resolution longitudinal data, substantiate and extent our previous cross-sectional baseline finding in the same cohort (Madsen et al., 2018). In general, neuroimaging studies rarely investigate regional asymmetry when investigating neuroticism and other negative emotionality-related traits. We previously observed associations between neuroticism and cingulum FA asymmetry in an independent cohort of healthy adults (Madsen et al., 2012). However, a recent study investigating associations between negative emotionality, assessed with the Child and Adolescent Dispositions Scale (CADS) at age 10-17 years, and cingulum FA asymmetry assessed 10-15 years later in a large cohort (n = 410), did not find any significant associations (Lahey et al., 2020). Inconsistencies between our and Lahey’s findings may in part be due to differences in personality questionnaires used, as the negative emotionality trait estimated with CADS exhibits only moderate correlations with neuroticism as well as weak, but significant correlations with conscientiousness and agreeableness (Lahey et al., 2020; Lahey et al., 2010).

Evidence supporting the importance of hemispheric asymmetry is more prominent in the electroencephalography (EEG) literature (Grimshaw & Carmel, 2014; Madsen et al., 2018; Nusslock et al., 2015). For example, two recent EEG studies observed that higher neuroticism scores were associated with increased left relative to right frontal alpha power during resting-state (Moshirian Farahi et al., 2019) or presentation of face stimuli with a direct gaze (Uusberg et al., 2015). Of note, alpha power is assumed to be inversely correlated with cortical activity, i.e., increased alpha power suggests lower neuronal activity (Allen et al., 2004; Nusslock et al., 2015). In the latter study, the neuroticism association was driven by facets related to avoidance behavior (i.e., anxiety, depression, self-consciousness and vulnerability), and not those related to approach behavior (i.e., angry hostility and impulsiveness). Interestingly, brain asymmetries have previously been described for approach/withdrawal behavior and emotional processing, with higher left relative to right frontal activity being associated with approach-related traits and positive affect, while lower left relative to right frontal activity has been linked to withdrawal-related traits and negative affect (Grimshaw & Carmel, 2014; Nusslock et al., 2015). Overall, the above suggests that cingulum asymmetry plays a role in neuroticism. It remains to be seen whether associations are driven by specific subcomponents of neuroticism. Furthermore, future studies should establish whether cingulum FA asymmetry plays a role in other negative emotionality traits.

We observed opposite relationships between neuroticism and cingulum FA asymmetry for females and males. Higher neuroticism scores were associated with increased left relative to right cingulum FA in females and decreased left relative to right cingulum FA asymmetry in males, substantiating our previous cross-sectional findings in the same cohort (Madsen et al., 2018). Since females and males did not significantly differ in cingulum FA asymmetry, the observed sex differences in neuroticism-cingulum associations are likely not driven by structural sex differences in cingulum FA per se. Opposite sex effects have also been observed in the relationship between neuroticism and grey matter volume and cortical thickness of the subgenual anterior cingulate cortex in adolescents aged 16-17 years, with females showing positive and males negative correlations (Blankstein et al., 2009). Further, a cross-sectional study in adults aged 19-80 years reported that higher neuroticism was associated with thicker anterior cingulate cortex in females and thinner anterior cingulate cortex in males with increasing age (Sweeney et al., 2019). Notably, the anterior cingulate cortex projects to and receives fibers from the cingulum. Overall, these findings suggest that there are sex differences in the associations between neuroticism and the cingulum and related cortical regions. However, the mechanisms underlying these associations are still unknown.

One possible explanation for the observed sex differences in neuroticism-cingulum associations may be differences in sex hormones. Supporting such notion are studies showing that differences in sex hormones may affect functional brain asymmetries (Hausmann, 2017). For instance, in an EEG study, frontal alpha asymmetry differed across the menstrual cycle in women depending on whether they had high or low neuroticism scores (Huang et al., 2015). Moreover, in a recent study in adolescent males, using transcranial Doppler ultrasonography, pubertal testosterone levels affected the extent of asymmetric brain activation during a mental rotation task as well as a chimeric face task (Beking et al., 2018). Intriguingly, the authors observed that the direction of the association depended on the levels of prenatal testosterone exposure, suggesting that prenatal and pubertal testosterone levels might affect functional brain asymmetries in a complex manner (Beking et al., 2018). All in all, these findings, make it plausible that sex hormones may play a significant role in the observed sex differences in neuroticism-cingulum associations, and thus warrants future study. Nevertheless, genetic differences, stress hormones and their interaction with sex hormones (see below), as well as psychological and/or socio-cultural factors may also be involved.

Our longitudinal study design allowed us to track changes in the neural correlates of neuroticism across time. In females, the relationship between neuroticism and cingulum FA asymmetry was already present in the youngest part of the included age range and appeared to be stable across childhood and adolescence. This suggests that the relationship arose earlier in life i.e., before late childhood, and we speculate that it might be innate. Indeed, (leftward) cingulum FA asymmetry appears to be present already in infants and remain throughout childhood, adolescence and early adulthood (Cohen et al., 2016). In line with this, we observed that (leftward) cingulum FA asymmetry did not significantly change with age, though FA increased significantly in both the left and the right cingulum. Furthermore, both neuroticism (Sanchez-Roige et al., 2018; van den Berg et al., 2014) and cingulum DTI measures (Budisavljevic et al., 2016; Peper et al., 2008; Shen et al., 2016; Vuoksimaa et al., 2017; Zhao et al., 2019) are moderately heritable, suggesting that at least part of the observed associations between neuroticism and cingulum FA asymmetry may be linked to genetic differences. Finally, since increased prenatal stress has been linked to increased fearfulness in toddlers (Bergman et al., 2007) and altered functional asymmetry in both rodents and healthy children (Alonso et al., 1991; Jones et al., 2011), as well as lower FA in the left cingulum in young adults (Marečková et al., 2019), we speculate that prenatal stress might also play a role in the observed associations between neuroticism and cingulum FA asymmetry.

Though we were unable to find statistically significant age-related changes in the link between neuroticism and cingulum FA asymmetry in males, visual inspection of the age trajectories of this relationship suggested that the association first became apparent during adolescence. Furthermore, our post hoc analyses of the youngest and oldest male participants suggested that the neuroticism-cingulum FA asymmetry relationship was stronger in the older age range. Speculatively, neuroticism-cingulum FA asymmetry associations may be less stable in males and may be influenced by moderating factors during adolescence. Interestingly, higher endogenous testosterone levels have been associated with lower neuroticism levels in adolescent and young adult males (Schutter et al., 2017). Furthermore, in the same study, lower neuroticism levels were linked to larger cerebellar volume, and this relationship appeared to be mediated by endogenous testosterone levels. As there is also evidence that pubertal testosterone may affect functional brain asymmetries in adolescent males, depending on the levels of prenatal testosterone exposure (Beking et al., 2018), we hypothesize that pubertal testosterone levels may affect the relationship between neuroticism and cingulum FA asymmetry during adolescence.

The current study has several strengths and limitations. A major strength of this study is its longitudinal design with high temporal resolution and up to 11 MRI scans per individual, optimized to investigate changes in brain-behavior relationships across late childhood and adolescence. Furthermore, since neuroticism scores did not significantly change with age in the investigated age range, we were able to use the mean neuroticism score, which allowed us to capitalize all the longitudinal MRI assessments as well as include participants with only one neuroticism assessment, thereby enhancing statistical power. Nevertheless, using mean neuroticism scores instead of longitudinal neuroticism scores might have prevented us from finding subtle changes in the relationship between neuroticism and associated white matter correlates. Additionally, we specifically covered the period of late childhood and adolescence (7-18 years), allowing us to trace changes in the peri-pubertal developmental period, where sex differences in neuroticism have been reported to emerge (De Bolle et al., 2015). However, since the cingulum show protracted maturation into early adulthood (Lebel et al., 2008), future studies should consider including a broader age range encompassing early childhood and adulthood. Finally, our sample size consisted of more females (n = 47) than males (n = 29), which might have prevented us from finding any significant neuroticism associations with cingulum FA asymmetry and neuroticism in males as well as significant sex differences in the association between neuroticism and cingulum FA asymmetry over age.

## 5. Conclusions

Using high temporal resolution longitudinal data, we show robust associations between trait neuroticism scores and cingulum FA asymmetry in typically-developing children and adolescents aged 7-18 years, with opposite effects in females and males. In females, higher neuroticism was associated with increased left relative to right cingulum FA, while in males, higher neuroticism was associated with decreased left relative to right cingulum FA. The association appeared to be stable across late childhood and adolescence in females, and we speculate that this association might be innate. In males, however, the association appeared to be stronger in the higher end of the investigated age range, and we hypothesize that increasing pubertal testosterone levels might play a role in this. Future studies should cover a larger age span to elucidate when the relationship between neuroticism and cingulum FA asymmetry arises, and to clarify the possible role of sex hormones, as well as genetic variation and prenatal stress exposure.

## Funding

This work was supported by the Danish council of Independent Research | Medical Sciences (grant numbers 09-060166 and 0602-02099B), the Lundbeck Foundation (grant number R32-A3161), the Lundbeck Foundation Center of Excellence grant to The Center for Integrated Molecular Brain Imaging, and EU Horizon 2020 research and innovation program grant for the Lifebrain project (grant agreement number 732592). Hartwig R. Siebner holds a 5-year professorship in precision medicine at the Faculty of Health Sciences and Medicine, University of Copenhagen, which is sponsored by the Lundbeck Foundation (grant number R186-2015-2138).

## Declaration of competing interest

Hartwig R. Siebner has received honoraria as speaker from Sanofi Genzyme, Denmark and Novartis, Denmark, as consultant from Sanofi Genzyme, Denmark, Lophora, Denmark, and Lundbeck AS, Denmark, and as editor-in-chief (Neuroimage Clinical) and senior editor (NeuroImage) from Elsevier Publishers, Amsterdam, The Netherlands. He has received royalties as book editor from Springer Publishers, Stuttgart, Germany and from Gyldendal Publishers, Copenhagen, Denmark. The authors declare no potential conflict of interest.

## CRediT authorship contribution statement

**Anna Plachti**: Conceptualization, Software, Formal analysis, Data curation, Writing - original draft, review and editing, Visualization. **William FC Baaré**: Conceptualization, Methodology, Software, Resources, Data curation, Writing - original draft, review and editing, Supervision, Project administration, Funding acquisition. **Louise Baruël Johansen**: Software, Formal analysis, Data curation, Writing - review and editing. **Wesley K. Thompson**: Software, Writing - review and editing. **Hartwig R. Siebner**: Resources, Writing - review and editing, Funding acquisition. **Kathrine Skak Madsen**: Conceptualization, Methodology, Software, Investigation, Resources, Data curation, Writing - original draft, review and editing, Visualization, Supervision, Project administration, Funding acquisition.

## Acknowledgements

**Acknowledgements**

The authors thank the children and their parents for their participation in the HUBU study.

## Abbreviations

DWI: diffusion weighted imaging
DT: diffusion tensor
DTI: diffusion tensor imaging
ROI: region-of-interest
FA: fractional anisotropy
AD: axial diffusivity
MD: mean diffusivity
RD: radial diffusivity
vmPFC: ventromedial prefrontal cortex
vmPFC_WM_: ventromedial prefrontal cortex white matter
UF: uncinate fasciculus
WM: white matter

