## Supplementary Material for "Only females show a stable association between neuroticism and microstructural asymmetry of the cingulum across childhood and adolescence: A longitudinal DTI study"

### Supplementary methods

#### Participants

Nineteen participants were excluded from further analyses, due to lack of acquired neuroticism data (n = 16), receiving a psychiatric diagnosis after study initiation (n = 2), or an incidental clinical finding on the MRI scan (n = 1). Our final sample consisted of 76 participants (47 females, 29 males) aged 7.5-18.9 years (mean = 12.4, standard deviation = 2.4). From these, we excluded 64 MRI sessions, because the participant did not finish the MRI scanning session (1 participant, 2 assessments), was not scanned due to metallic dental braces (13 participants, 30 assessments), had poor MR-image quality (17 participants, 23 assessments), had acquired a brain injury after baseline (1 participant, 8 assessments) or the assessment was accidently left out from the preprocessing (1 assessment).

#### Image acquisition

"All subjects were scanned using a 3T Siemens Magnetom Trio MR scanner (Siemens, Erlangen, Germany) with an eight-channel head coil (Invivo, FL, USA). All acquired scans were aligned parallel to the anterior commissure–posterior commissure line. T1-weighted images of the whole head were acquired using a 3D MPRAGE sequence (TR = 1550 ms, TE = 3.04 ms, matrix 256 x 256, 192 sagittal slices, 1 x 1 x 1 $mm^3$ voxels, acquisition time = 6:38). T2-weighted images of the whole head were acquired using a 3D turbo spin echo sequence (TR = 3000 ms, TE = 354 ms, FOV = 282 x 216, matrix = 256 x 196, 192 sagittal slices, 1.1 x 1.1 x 1.1 $mm^3$ voxels, acquisition time = 8:29). Whole brain diffusion-weighted (DW) images were acquired using a twice-refocused balanced spin echo sequence that minimized eddy current distortion (Reese et al., 2003). Ten non-DW images (b = 0) and 61 DW images (b = 1200 $s/mm^2$), encoded along independent collinear diffusion gradient orientations, were acquired (TR = 8200 ms, TE = 100 ms, FOV = 220 x 220, matrix = 96 x 96, GRAPPA: factor = 2, 48 lines, 61 transverse slices with no gap, 2.3 x 2.3 x 2.3 $mm^3$ voxels, acquisition time = 9:50). A gradient echo field map was acquired to correct $B_0$ field distortions (TR = 530 ms, TE[1] = 5.19 ms and TE[2] = 7.65 ms, FOV = 256 x 256; matrix = 128 x 128, 47 transverse slices with no gap, voxel size = 2 x 2 x 3 $mm^3$, acquisition time = 2:18)." (Madsen et al., 2018).

#### Tract-based spatial statistics

"Tract-Based Spatial Statistics (TBSS) (Smith et al., 2006), part of FSL 5.0.9, was used to create a mean FA skeleton, representing the centers of all tracts common to the group. Instead of using the standard TBSS normalization steps, the between-subject FA template image (from DTI-TK) was aligned to MNI space using affine registration (flirt, FSL). Subsequently, the between-subject diffusivity images and the normalized individual DT images were transformed into 1 $mm^3$ MNI space. Next, the MNI space aligned, between-subject FA template image was entered into the TBSS processing stream using the "tbss_skeleton" script, part of the "tbss_3_postreg" processing step, in which the between-subject FA template image was thinned to create a mean FA skeleton. The mean FA skeleton was thresholded at FA > 0.2 and contained 102,983 $1mm^3$ interpolated isotopic voxels, corresponding to approximately 22% of the voxels (in the mean FA map across subjects and time points) with FA above 0.2." (Madsen et al., 2020). Next, all participants' aligned FA images were projected onto the mean FA skeleton by locating the voxels with the highest local FA value perpendicular to the skeleton tracts and assigning these values to the skeleton. Finally, the skeleton projections were applied on the AD and RD data.

#### Regions-of-interest

White matter ROIs included the cingulum, the uncinate fasciculus (UF) and the white matter underlying the ventromedial prefrontal cortex ($vmPFC_{WM}$). The cingulum ROIs included all skeleton segments within the cingulum, and excluded any segments intersecting but diverging from the main body of the cingulum. The skeleton segments representing the cingulum were clearly distinguishable from all other skeleton segments. The left and right cingulum ROIs contained 643 and 622 voxels, respectively. The $vmPFC_{WM}$ ROIs included the skeleton segments in the white matter underlying the left and right vmPFC, while excluding segments in the frontal pole. The $vmPFC_{WM}$ ROIs extended between MNI-coordinates y = 51 to y = 33, both included. The left and right $vmPFC_{WM}$ ROIs contained 142 and 162 voxels, respectively. The UF ROIs were delineated using the JHU White-Matter Tractography Atlas (Hua et al., 2008) implemented in FSLview for guidance. The ROIs included central UF segments, while skeleton segments extending towards the inferior frontal gyrus, the orbitofrontal cortex and the temporal pole were excluded. The left and right UF ROIs included 335 and 356 voxels, respectively. Finally, to test the anatomical specificity of observed associations, left (50,795 voxels) and right (51,843 voxels) hemispheric ROIs, including all skeleton segments within each hemisphere, were delineated using the mid-sagittal plane (not included in either of hemispheric ROIs).

### Supplementary results

#### Effect size maps

The effect size maps display the t-values where FA was positively (red-yellow colors) or negatively (blue-light blue colors) associated with $neuroticism_{mean}$, controlling for age and RMS. The full unthresholded t-maps can be downloaded from https://neurovault.org/collections/LRMGYBHV/. In the description below, the numbers between brackets represent the MNI Y coordinate depicted in Supplementary Figure 1.

In females, $neuroticism_{mean}$ was positively and negatively associated with FA ($3.3 \leq t \leq -3.3$, $p < 0.001$, uncorrected), in respectively, 1.17% and 4.12% of all skeleton voxels, excluding voxels for which the GAMM model did not converge (0.44%). Higher $neuroticism_{mean}$ scores were associated with higher

FA in several clusters across the white matter skeleton, including the splenium of corpus callosum (-43) and the temporal part of the right superior longitudinal fasciculus (-50), as well as the white matter underlying the left anterior parahippocampal gyrus (2), and left supramarginal gyrus (-50). Furthermore, lower $neuroticism_{mean}$ was associated with higher FA in multiple white matter regions, including the left inferior fronto-occipital fasciculus (33), right UF (8), left inferior longitudinal fasciculus (-9), and bilateral anterior thalamic radiation (-9) as well as the white matter underlying the middle and superior frontal gyrus (24), right temporal pole (8), right inferior precentral gyrus (-3), left subcentral gyrus (-3, -6), left superior temporal gyrus (-16), right supramarginal gyrus (-36), and right middle temporal gyrus (-40).

In males, we found that $neuroticism_{mean}$ was positively and negatively associated with FA in, respectively, 4.2% and 2.3% of all skeleton voxels, excluding voxels for which the GAMM model did not converge (0.34%). Higher $neuroticism_{mean}$ was associated with higher FA in multiple clusters across the white matter skeleton, including the right UF (43, 2), genu of corpus callosum (26), bilateral inferior fronto-occipital fasciculus (43, -49, -73), right corticospinal tract (-30), bilateral superior longitudinal fasciculus (-30, -49), left posterior corona radiata (-35), bilateral posterior thalamic radiation (-49, -51), bilateral inferior longitudinal fasciculus (-49, -73), forceps major (-73) as well as the white matter underlying the left superior frontal gyrus (43), bilateral inferior frontal gyrus (26, -3), right supplementary motor area (-3), bilateral postcentral gyrus (-18), and right superior parietal lobule (-81). Further, lower $neuroticism_{mean}$ scores were associated with higher FA in right UF/inferior longitudinal fasciculus (2) and right forceps major (-81) as well as the white matter underlying the left superior frontal gyrus (37), left posterior frontal opercular cortex (2), and left precuneus (-49).

Additional t- and F-maps of the association of FA with $neuroticism_{mean}$-by-age for the whole group and females and males separately have also been uploaded to NeuroVault.

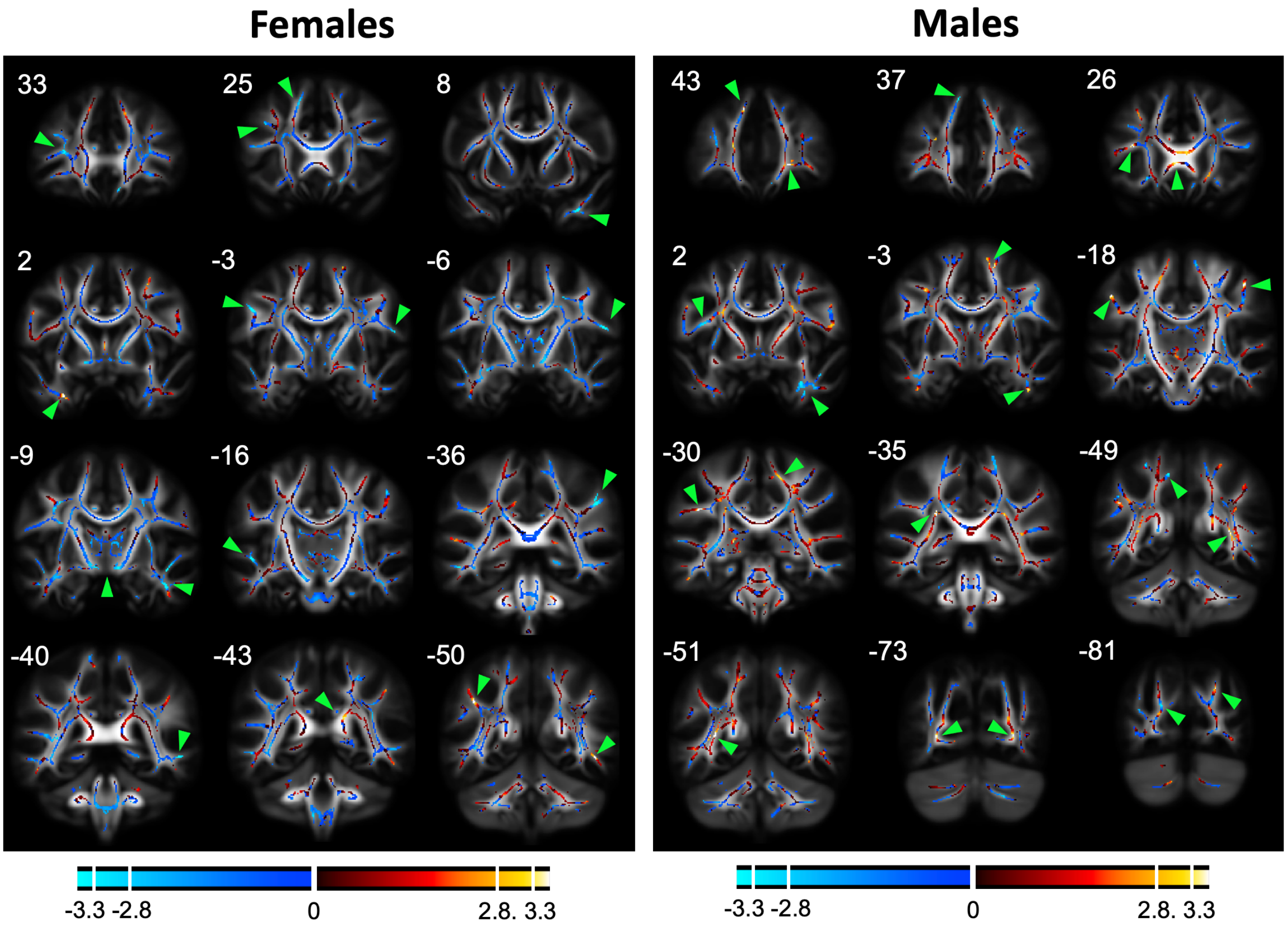


**Supplementary Figure 1**. Effect size map displaying the association between $neuroticism_{mean}$ and FA across the white matter skeleton, corrected for age and RMS. The effect size map displays the images with the highest t-values and largest clusters, based on visual inspection, with the main clusters pointed out using green arrows. Voxels, where higher $neuroticism_{mean}$ was associated with higher FA, are depicted in warm colors ranging from red to yellow, while cold colors ranging from dark blue to light blue depict where higher $neuroticism_{mean}$ was associated with lower FA. The t-values in the color bar correspond to t = ± 3.3, p = 0.001 and t = ± 2.8, p = 0.005 (df = 366 for females, df = 226 for males, two-tailed, uncorrected). The MNI Y coordinates for the coronal slices are given above each image. Images are shown according to neurological convention, with the left hemisphere depicted in the left side.
